# Expansion stimulated emission depletion microscopy (ExSTED)

**DOI:** 10.1101/278937

**Authors:** Mengfei Gao, Riccardo Maraspini, Oliver Beutel, Amin Zehtabian, Britta Eickholt, Alf Honigmann, Helge Ewers

**Affiliations:** Institut für Biochemie, Freie Universität Berlin, Thielallee 63, 14195 Berlin, Germany; Max Planck Institute of Molecular Cell Biology and Genetics, Pfotenhauerstraße 108, 01307 Dresden, Germany; Institut für Biochemie, Charité Universitätsmedizin Berlin

## Abstract

Stimulated emission depletion (STED) microscopy is routinely used to resolve the ultra-structure of cells with a ∼10-fold higher resolution compared to diffraction limited imaging. While STED microscopy is based on preparing the excited state of fluorescent probes with light, the recently developed expansion microscopy (ExM) provides sub-diffraction resolution by physically enlarging the sample before microscopy. Expansion of fixed cells by crosslinking and swelling of hydrogels easily enlarges the sample ∼4-fold and hence increases the effective optical resolution by this factor. To overcome the current limits of these complimentary approaches, we here combined ExM with STED (ExSTED) and demonstrate an increase in resolution of up to 30-fold compared to conventional microscopy (<10 nm lateral and ∼50 nm isotropic). While the increase in resolution is straight forward, we found that high fidelity labelling via multi-epitopes is required to obtain emitter densities that allow to resolve ultra-structural details with ExSTED. Our work provides a robust template for super resolution microscopy of entire cells in the ten nanometer range.

## Introduction

Expansion microscopy (ExM), which allows super-resolution imaging with conventional microscopy systems has been proposed in 2015^1^. In this method, immunofluorescence stained cells or tissues are physically enlarged by a procedure that starts with embedding a fixed and immunolabelled cell into a polyacrylamide gel. Immunofluorescence probes are then crosslinked to the gel matrix and the original sample is degraded by proteases. In this way, the position of fluorophores is stored directly in the gel matrix. The hydrogel is then swollen by exchanging the buffer for deionized water. As a result, the distance between the fluorescent probes crosslinked to the gel is increased, which allows to resolve previously optically inaccessible distances and effectively a higher spatial resolution.

ExM is developing at a high pace^2–7^, by now it allows for conventional immunofluorescence labelling approaches^6^ and it has been tested on different samples from pathological preparations^3^ to brain sections^4^. Higher expansion factors up to 10-fold or even 20-fold have been reported using a different polymer^8^ or a two-step expansion protocol^2^. However, it remains unclear, to what extent the expanded cellular structures truly represent the native conformation in cells below the resolution limit. It was recently shown that organized arrays of tightly spaced microtubules in the protozoan pathogen *Giardia lamblia* remain ordered after ExM with subsequent structured illumination microscopy (SIM)^9^, demonstrating convincing isotropic expansion at a high resolution level. Nevertheless, the resolution stopped short of what can be routinely achieved in single molecule localization microscopy (SMLM).

In principle, combining ExM with diffraction unlimited super-resolution techniques like PALM/STORM^10–12^, stimulated emission depletion microscopy (STED)^13^ or MINFLUX^14^ should enable a spatial resolution down to the scale of single proteins (<10 nm). However, the formation of a spatially expanded hydrogel during ExM introduces some technical challenges for super-resolution microscopy. First, because the sample is expanded, whole cell imaging requires to focus four times deeper into the sample compared to conventional samples. Therefore, water immersion lenses with a lower collection efficiency instead of oil-immersion have to be used to prevent spherical aberrations. Second, introduction of special buffers required for stochastic on and off switching of fluorophores^1,12^ in SMLM are likely to interfere with the expanded gel because of osmotic imbalances. Third, the labelling density required for meaningful <10 nm imaging needs to be very high, which is currently difficult to achieve with ExM because a significant fraction of the dyes is lost due to the digestion of antibodies during the preparation procedure.

Keeping these limitations in mind, we decided to combine ExM with STED. First STED is based on a confocal setup, therefore imaging deep in the sample is relatively straight forward and secondly STED does not require any special buffers. Finally, to overcome the labelling density problem we used a brute force multi-epitope labelling approach.

We report here an ExSTED imaging protocol that allows for below 10 nm two-dimensional (2D) resolution and isotropic 50-70 nm three-dimensional (3D) resolution in fixed cells.

## Results

We first aimed to optimize resolution and contrast in ExM towards the routine imaging of nanoscopic structures in cells. We immunostained microtubules in fixed cells, polymerized and expanded a hydrogel in the sample. When we then imaged the gels in a spinning disc confocal microscope, we found that expansion was isotropic and yielded significantly higher resolution after considering the expansion factor (Figure 1a – c, Figure S1a-c). However, we noticed a strong loss of fluorescence signal, even when we used directly labelled primary antibodies (Figure S1a and ^2–7^). This is not surprising, given that the usually achieved expansion factor of ∼4 in a single dimension translates into a factor of 64 in the volume and thus a proportionally lower signal in a given area. To overcome this problem, we developed an intensive labelling method by expressing α-tubulin-GFP in HeLa cells and then using a cocktail of anti-GFP, anti-α-tubulin and anti-β-tubulin antibodies for immunofluorescence staining to cover a maximal number of epitopes per microtubule. Each antibody worked efficiently in conventional microscopy (data not shown). We then applied AF488-labelled secondary antibodies against all primary antibodies. We found that this ultra-dense labelling approach resulted in significantly brighter (∼4x) microtubule staining after expansion of the gel (Figure 1d-f, Figure S1d-h, Figure S2). We concluded that intense multi-epitope labelling allows for excellent signal to noise ratio in ExM.

**Figure 1:**
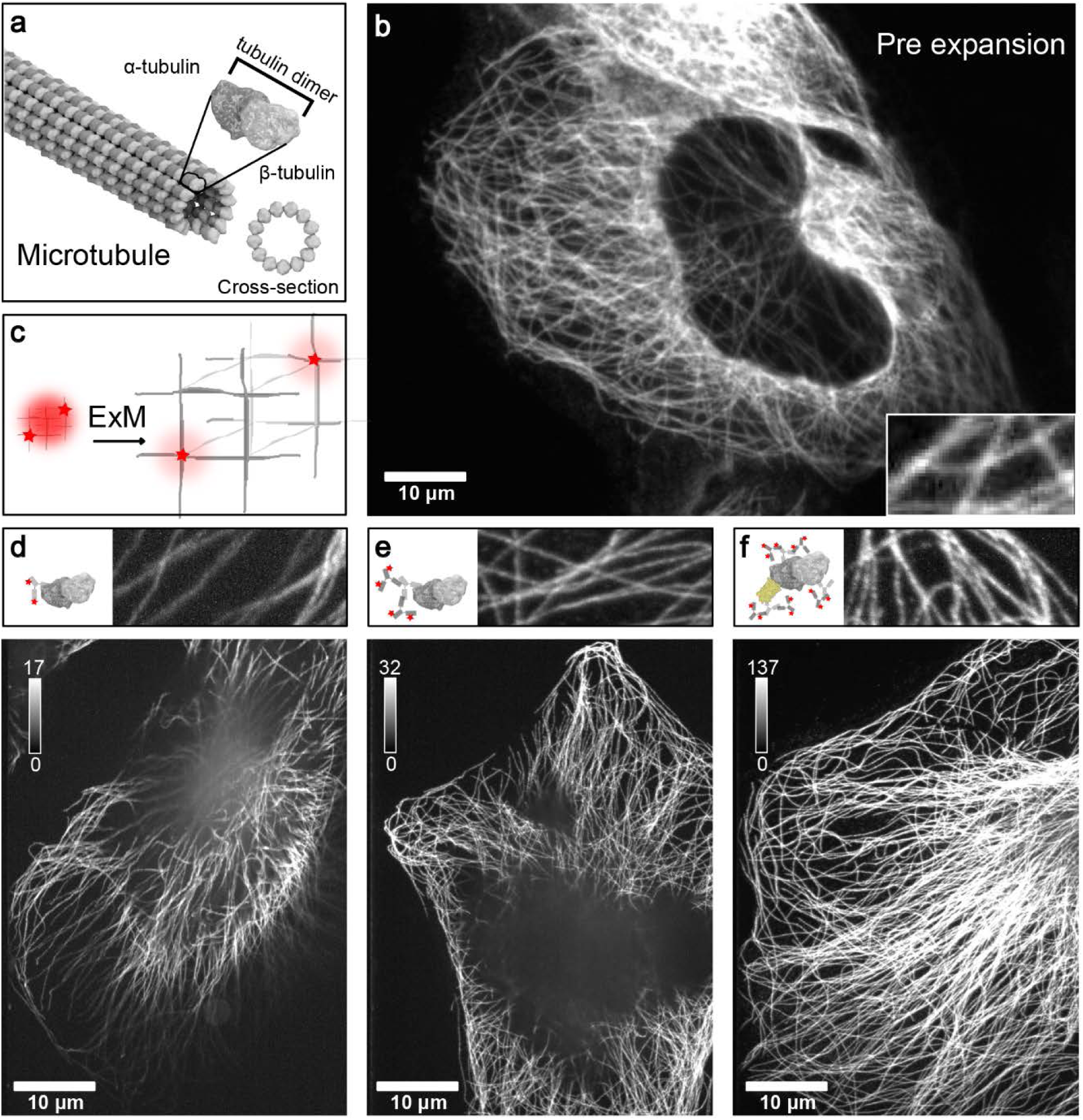
Improved labelling for ExM. **(a)** Illustration of the ultra-structure of a microtubule. **(b)** Hela cell stained with anti-α-tubulin antibody. Inset is enlarged 4 times. **(c)** Schematic of the expansion process. The red stars represent two dye molecules, the light red circle represents the diffraction limit of light. After ExM, the distance between the dye molecules is increased due to the expansion of the hydrogel (grey matrix), so that the two dyes can be resolved individually. However, the brightness of each signal is half of the combined signal before expansion. **(d – f)** Microtubules in HeLa cells immunostained by different protocols and imaged after the ExM treatment. Shown are maximum projections of 3D fluorescence micrographs with different intensity scaling demonstrating a ∼4fold increase of signal using the multi-epitope labelling. The top images are magnified 4 times from the bottom images. **(d)** Hela cell stained only with AF488 labelled anti-α-tubulin antibody. **(e)** Hela cell stained with anti-α-tubulin antibody and an AF488 labelled secondary antibody. **(f)** Hela overexpressing α-tubulin-GFP after staining with anti-α-tubulin, anti-β-tubulin and anti-GFP-primary antibodies and AF488 labelled secondary antibody. All images were taken by spinning disk confocal microscopy. All distances and scale bars were calculated with the expansion factor. Scale bar is 10 µm.

We hypothesized that multi-epitope labelling would allow for sufficient signal to perform STED microscopy of expanded gels and thus combine the factor 4 of expansion with the increased resolution of STED microscopy. To realize this experiment, we performed multi-epitope labelling on microtubules using secondary antibodies coupled to the dye Abberior Star Red, which is optimized for STED microscopy. Our STED-microscope allows for a lateral resolution of 30 nm in 2D mode and an axial resolution of ∼100 nm in 3D mode with a high NA objective after alignment. Using our STED setup, we then scanned expanded gels in 2D mode to visualize individual microtubules. We found that in ExSTED images, microtubules appeared as pairs of aligned, mostly continuous structures as reported in highest resolution single molecule localization microscopy methods^6,15–17^ (Figure 2a, b). The two lines were separated by 38.6 ± 11.9 nm (Figure 2c), consistent with an average displacement of the dye from the microtubule surface of ∼ 7 nm due to the antibodies^3,18^ as has been found in previous SMLM measurements^2,4,19^. To determine the effective resolution after expansion and STED imaging, we fitted single spots with a 2D Gaussian and line-profiles through microtubules using a 1D Gaussian. Both approaches gave a resolution in XY of 8 ± 1.4 nm (full width at half maximum) (Figure 2c). This demonstrates an outstanding resolution which approaches the size of the label itself (primary plus secondary antibody). Additionally, it shows that the native microtubule structure is represented faithfully by the expanded gel down to the nanoscopic range. We concluded that ExSTED can resolve cellular structures less than 10 nm apart.

**Figure 2:**
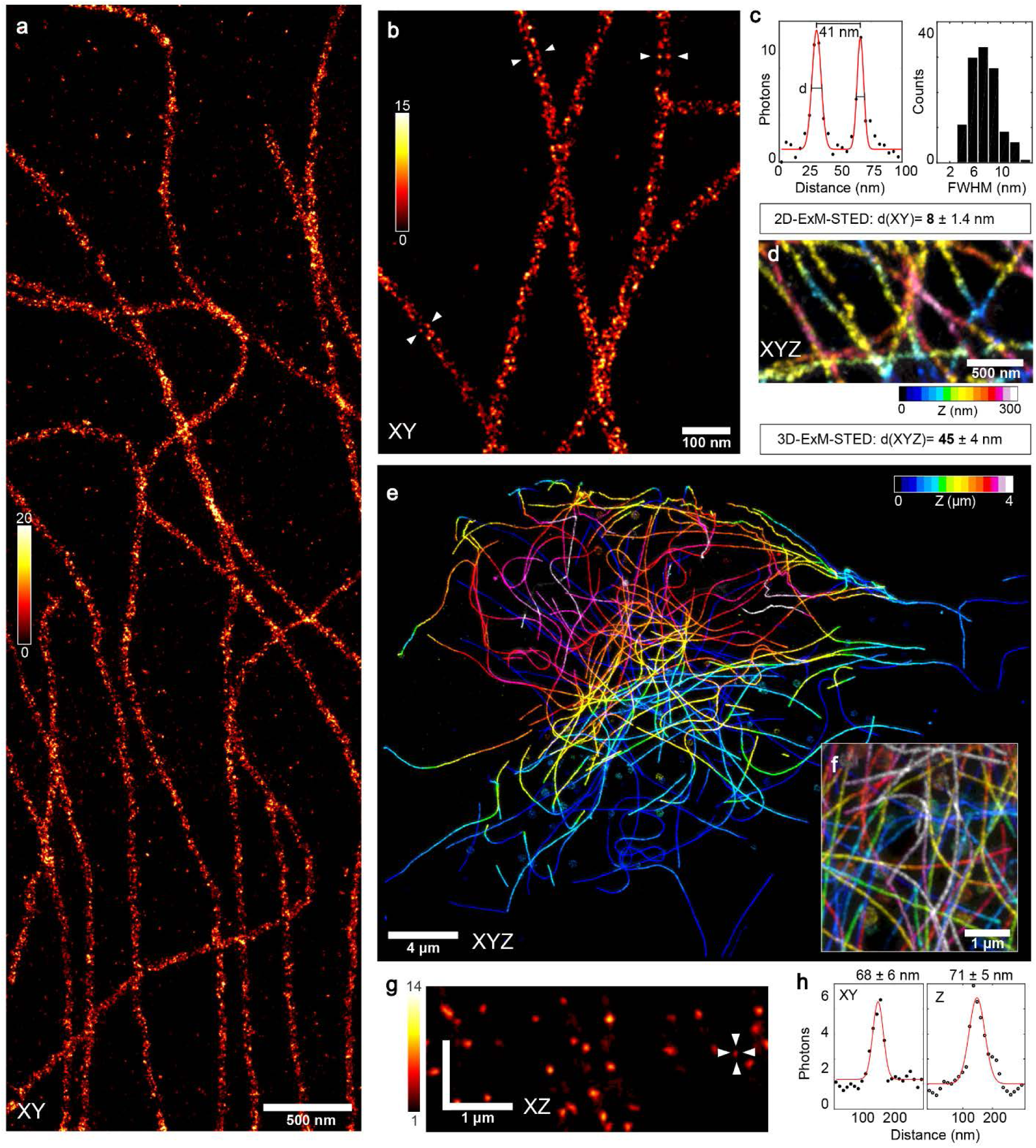
Combination of ExM with STED. **(a)** 2D-STED of expanded tubulin staining using an 100x oil immersion lens imaging close to the cover glass surface. **(b)** Magnified view of the microtubules. Note the resolution allows to clearly distinct antibody positions on opposing sites of a single tubule. **(c)** Left panel: Quantification of the apparent diameter of microtubules including primary and secondary antibodies. Right panel: Quantification of the 2D resolution in image **(a)**. **(d)** 3D-STED of expanded tubulin staining using the oil objective close to the surface. The Z position is indicated by the color coding. **(e)** Whole cell 3D-ExSTED using a water 60x immersion lens to enable greater depth penetration. **(f)** Close up of a dense region in 3D. **(g)** XZ cross-section of from **(f)**, note the isotropic shape of the PSF. **(h)** Exemplary XY and Z profiles indication an isotropic resolution of ∼ 70 nm. Scale bar is 500 nm in **(a)**, 100 nm in **(b)**, 500 nm in **(d)**, 4 µm in **(e)** and 1 µm in **(f)** and **(g)**.

We next aimed to test, if we could achieve isotropic sub 100 nm resolution in cells using ExSTED microscopy. To do so, we employed the 3D mode of our STED microscope. First, we used the oil immersion lens to achieve the highest possible resolution close to the surface (< 5 µm). Figure 2d shows a 3D XYZ image of a dense microtubule staining. In this configuration, we achieved an isotropic resolution of 45 ± 4 nm, which is very close to the diameter of microtubules determined with 2D ExSTED (Figure 1). This shows that the expansion protocol indeed produced an isotopically expanded sample down to nanometer range. We verified negligible distortion by the expansion (< 1 %) using a global analysis (Figure S1c). Since the gel consists mainly of water and has a RI of ∼ 1.33, whole cell imaging with the oil objective was impossible due to spherical aberration at depth above 5 µm. To overcome this problem, we used a 1.2 NA 60x water objective. This configuration slightly decreased isotropic resolution to 70 nm but we were able to resolve the complete microtubule network with excellent signal to noise (Figure 2e-h). Because scanning such large field of views in 3D with nanometer pixel sizes takes hours, we had to minimize the drift of the sample induced by loss of water from the gel or other interactions. To prevent drift, we completely embedded the cover glass mounted gels into two component silicone. This greatly reduced drift and allowed imaging of the same sample up to days. The remaining drift was corrected for using cross-correlation between the slices of the image stacks. The 3D images acquired with this procedure allowed to resolve close crossings of microtubules in dense areas (Figure 2e, f, Supplementary MovieS1). We concluded that ExSTED allowed for isotropic imaging of entire cells in 3D with 70 nm resolution.

While the accurate measurements of subresolution distances between giant amorphous multiprotein complexes such as the pre-and postsynapse or within clathrin-coated pits have been repeatedly reported in ExM^1,6,8^, it remains unclear, whether the organization and orientation of multiprotein structures remains intact at the nanoscopic level in ExM. To investigate if this is possible in principle, we investigated a number of structures, whose organization is not accessible in conventional light microscopy, but whose size is well established by super-resolution and electron microscopy methods. Specifically, we investigated the periodic actin-spectrin ring structure found along neuronal axons^2,20^, centrioles^9,21^, primary cilia of mammalian cells^10–12,22^ and motile cilia^13,23^.

Since we were able to resolve closely spaced microtubules in 3D, we next characterized a structure containing 9 microtubule doublets, the primary cilium in Madin-Darby canine kidney (MDCK) epithelial cells (Figure 3a, b). We immunostained confluent layers of MDCK cells against acetylated tubulin and the cilia membrane protein Arl13b. The primary cilium grew from the top of the cells and projected upwards, rendering access by light microscopy with oil immersion difficult due to loss in signal and optical aberrations. We decided to employ three different approaches towards overcoming this problem. i) we grew MDCK monolayers and then used poly-L-lysine coated coverslips to bind to the cilia and rip them off the apical side of the monolayer. In this configuration the cilia are oriented parallel to the cover glass surface and therefore resolving the axial symmetry of the 9 doublets is difficult ii) We used a water immersion lens to access cilia on the apical side with improved working distance of the objective and less aberrations, but with reduced resolution. iii) We tried to match the refractive index of the sample^14,24,25^ towards the oil immersion using sucrose solution to reduce optical aberrations. Iodixanol^25^ lead to shrinking of the gel and could not be used. Soaking the gel in a highly-concentrated sucrose solution (80% w/v) however lead to an acceptable degree of shrinkage of the expanded gel (10%). We found that refractive index matching for the oil objective or using water immersion objectives yielded similar resolution in 2D ExSTED images of cilia. When we imaged cilia using ExSTED, we found that a coat of Arl13b staining surrounded a core of acetylated tubulin as expected (Figure 3c). At the basal parts of the cilia we were able to resolve the microtubule doublets inside the cilium (Figure 3d) surrounded by a thin layer of Arl13b as (Figure 3e), consistent with a membrane staining around a cilium (see Figure 3a). We then turned to a cilia structure that contained a central microtubule, the *Chlamydomonas reinhardtii* motile cilium (Figure 3f). Using ExSTED we resolved the 9 + 2 microtubules in cross-sections of the cilia (Figure 3g) and identified individual microtubule doublets along the length of the cilium (Figure 3h). Last, we tried to image centrioles in HeLa cells using immunofluorescence staining against CEP152 (see Figure 3i). Using ExSTED, we could clearly resolve pairs of centrioles as ring-shaped structures (top-view) or hollow tubes (side view). Additionally, we found that CEP152 was enriched on one side of the centriole, which is consistent with previous studies describing CEP152 on the proximal side of the centriole (Figure 3j). Similarly, we also were able to resolve the 190-nm spaced ring structure around axons in primary mouse hippocampal neurons (Figure S3). We concluded that the use of index matching in ExSTED allowed for optimal resolution at depths of up to 50 μm in the expanded sample that are compatible with whole cell 3D imaging at a resolution of tens of nanometers.

**Figure 3:**
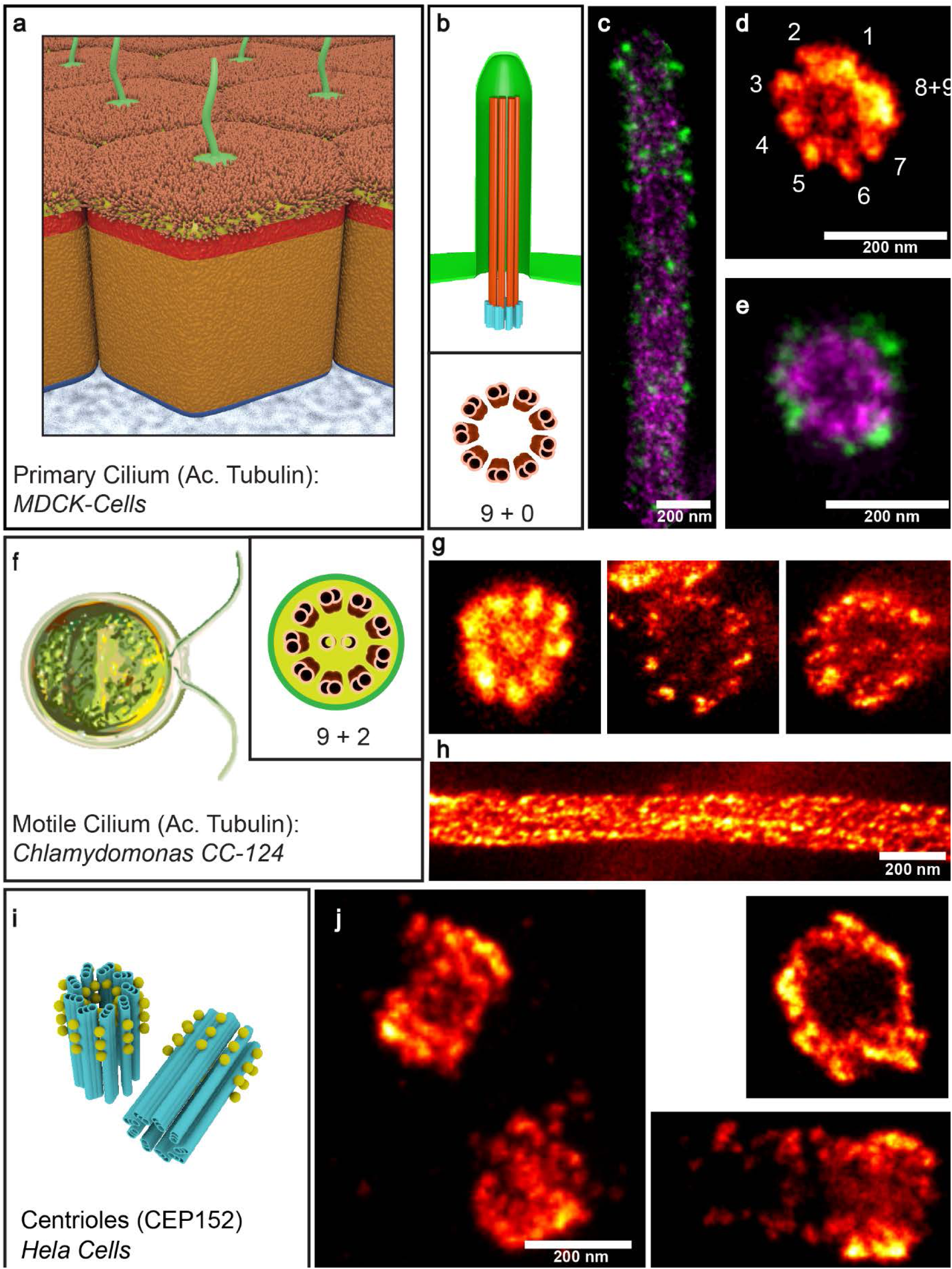
ExSTED of primary and motile cilia and centrioles. **(a)** Model of mammalian epithelial MDCK cells with primary cilia at the apical side. **(b)** Model of the primary cilium showing the 9 microtubule doublets, with acetylated (ac.) tubulin in orange and the cilia membrane protein Arl13b in green. Lower panel shows a model of the cross section through the cilium. **(c)** Two color ExSTED on ac. tubulin (magenta) and Arl13b (green) showing a cilium cut on the long axis. **(d)** Single color image of ac. tubulin in cross section view. **(e)** Two color image as in **(c)** in cross section view. **(f)** Model of the motile cilia from Chlamydomonas reinhardtii showing the 9 microtubule doublets with the central +2. **(g)** Cross-sectional views of ac. tubulin of motile cilia at different depth in the sample. At increased depth spatial resolution is decaying due to spherical aberrations. **(h)** Cilium as in **(g)** in longitudinal view. **(i)** Model of the centrioles (blue) and the centriole associated protein CEP152. **(j)** ExSTED of CEP152 staining in Hela cells in different orientations. Note the asymmetric distribution of CEP152 towards one side of the centriole. All scale bars 200 nm.

## Discussion

Here we labelled cellular structures with high density to allow for ultra-high spatial resolution imaging using a combination of ExM with super-resolution STED microscopy. A four-fold expansion of the sample followed by STED microscopy of fluorescent dyes in the expanded sample enabled an effective spatial resolution below 10 nm in 2D and in the range of 50-70 nm in 3D.

In recent work, ExM has been combined with SIM^9^ yielding very low distortion images at super-resolution level. In this work, the authors demonstrated the resolution of individual microtubule doublets in flagella in close proximity to the coverslip. We found that ExSTED allows for similar resolution deep inside the sample (up to 50 µm) and for ultra-high resolution (<10 nm) if structures are within few µm above the coverslip. The imaging of microtubules as parallel lines around 40 nm apart we achieved here is a benchmark for high quality SMLM super-resolution imaging^15–17,19,26^. Similarly, the hollow nature of spectrin rings in neuronal axons has so far only been reported for 3D-SMLM^20,27,28^. ExSTED thus provides at least similar resolution to the most powerful super-resolution approaches available at the time, both in 2D and in 3D.

The possibility of distortions from the native arrangement of moleculees in cells through the expansion process is an important problem that must be addressed in the establishment of the technique. We find here that microtubules expanded isotopically, spectrin rings around axons are as circular as expected and the organization of microtubule doublets in cilia and the circumference of the cilia by closely-apposed membrane-associated molecules remains intact after expansion and STED imaging. While this does not rule out that other cellular structures deviate from the native structure in expanded samples, and occasionally samples are mechanically disrupted on a global scale through uneven digestion, such artifacts are easily spotted. In our samples, we find no evidence for sudden ruptures in individual sub-resolution structures that would not be present in comparable SMLM experiments in cells.

Although we established a labelling protocol that allowed for intense staining of target epitopes, this can be further improved upon. An ideal candidate is DNA-based technology^29^, as DNA is not degraded during the ExM preparation and it can be used to specifically deliver the fluorescent probe of choice into the gel^1^. Another factor is the spatially accurate delivery of the dye to the target, which is especially important in super-resolution microscopy as the resolution approaches the size of the labelling agent. Here, nanobodies may provide a solution^30,31^, however in our hands, tubulin or GFP nanobodies were not useful as labelling reagents for ExM and lead to high loss of fluorescence during the preparation. This was likely because they became degraded in the ExM procedure and the dye-coupled nanobody peptide was not crosslinked to the gel.

The expansion of entire cells creates facsimiles of single cells that are tens of micrometers thick, which is challenging for a high and super-resolution techniques. This fact will be even more significant for protocols that lead to higher expansion factors^2,8^. To address this problem, we explored refractive index matching of the hydrogel suitable for high NA objectives (silicone and oil). We found that sucrose is one of the lowest cost chemicals that allows a big improvement on the sample. In sucrose solution, Abberior star RED and AlexaFluor 594 didn’t show significant difference in brightness or bleaching speed. By losing 10% of the expansion factor, we were able to expand the usable working distance for oil immersion objectives up to 20 μm above the surface. However, using recently introduced silicone immersion (RI = 1.4) objectives should improve contrast and resolution of 3D ExSTED at depth of >20 μm even more, since with sucrose embedding an RI of 1.4 can be perfectly matched.

In general, the parameters that can be tuned in ExM preparations to achieve optimal resolution in super-resolution experiments are not yet fully explored. While we optimized the protocol for brightest staining and highest resolution in ExSTED, the procedure may be adapted to achieve optimal results under other experimental conditions that require changing the expansion factor and gel handling or the optical properties of the gel.

In summary, we have here introduced an approach to further improve the resolution of STED microscopy in 3D by combining it with ExM. We have demonstrated the possibility to resolve sub-10 nm structures in 2D and sub-50 nm structures in 3D. Our work thus provides access to a resolution previously only accessible with the best SMLM and opens the door to further advances using a combination of super-resolution and ExM techniques.

## Material and Methods

### Cell culture, fixation and staining

Primary hippocampal neurons were cultured and immunostained as described before^32^. In short, primary mouse hippocampal neurons were plated on Poly-L-lysine coated coverslips, grown for 15 days *in vitro* (DIV) and fixed with 4% paraformaldehyde (PFA) plus 4% sucrose in PBS. The neurons were subsequently permeabilized with 0.2% Triton X-100 in PBS and then blocked first with Image-IT FX (Invitrogen) and then with blocking buffer (3% BSA/0.1% Triton X-100 in PBS). Antibodies were diluted in the blocking buffer and incubated with the coverslips at room temperature for 1 h.

Hela and U2OS cells were cultured in medium with 10% FBS and 1% Glutamax in DMEM (all Life Technologies) and then transfected with Neon^®^ transfection system (ThermoFisher) according to the manufacturer’s instructions, using a plasmid encoding for pEGFP-α-tubulin (Clontech Cat. No. 632349). The transfected cells were cultured on 12 mm #1.5 coverslips (Thomas Scientific) in a 24-well plate for 48-72 h in the same growth medium. Cells were washed intensively with PBS and subsequently fixed. Cells shown in Figure 1 were fixed with −20 °C cold methanol for 5 min. And Figure 2 was achieved by 60 s of permeablization with 0.2 % TritonX-100, 1 mM EDTA in BRB80 (80 mM PIPES, 1 mM MgCl_2_ and 1 mM EGTA) and then 10 min incubation of 4 % PFA (ThermoFisher), 0.1 % glutaraldehyde (Electron Microscopy Sciences) in BRB80. Both samples were intensively washed after fixation, and blocked with 3 % BSA, 0.1 % Triton-X100 in PBS for 1 h at RT then stained with antibodies in the same buffer.

MDCKII cells were cultured for 2 weeks in MEM supplemented with 110 mg/L sodium pyruvate, 2 mM NEAA and 5 % FBS and fixed with 4 % PFA in PBS. The fixed cells were washed intensively with PBS, then blocked and stained in the same buffer as Hela.

### Gelation, digestion and expansion

Monomer solution (1 M NaCl, 8.625% sodium acrylate, 2.5% or 20% acrylamide and 0.15% N,N’-methylenebisacrylamide in PBS) was prepared in aliquots, stored at −20 °C, and thawed every time before use. The stained coverslips were first incubated 12 h in 0.1 mg/ml Acryloyl-X, SE (ThermoFisher) at room temperature, then washed with PBS for 3 times, finally changed to the monomer solution and incubated for 10 min. Hydrogels were formed by adding a final concentration of 0.15 % of tetramethylethylenediamine and 0.15 % ammonium persulfate to the monomer solution and putting the coverslip upside down to the drop of solution. Gelation took place in a humidity incubator at room temperature for 2 h. Proteinase K (New England Biolabs) was diluted to 8 U/ml in the digestion buffer (50 mM Tris-HCl, 1 mM EDTA, 0.1 % TritonX-100, 0.8 M guanidine HCl, pH 8.0) and applied to the sample in a 6-well plate at 37 °C for at least 3 h. Digested gels were transferred to dishes with MilliQ water for further expansion. Water was exchanged every 10 min for 3 - 5 times until the gel spread out in all dimension. 84%(w/v) sucrose solution was prepared freshly every time and added to the expanded gel at room temperature overnight. The gels were gently dipped onto a paper tissue for several times and mounted to a coverslip. The coverslip for imaging was first plasma cleaned, then incubated with poly-L-lysine and finally air dried after removal of liquid. Gels that were imaged for more than 1 h were further fixed using picodent twinsil®.

The expansion factor was calculated by overlapping the pre-and post-expanded gel air-water boundary or by the distances between landmarks in the pre-and post-cells. Thickness of the gel was calculated from the volume of the gelation solution and the surface area for pre-expanded gel, and measured by Vernier caliper for the post-expanded gel.

### Microscope and image analysis

Standard immunofluorescence microscopy was performed on an inverted Olympus IX71 microscope equipped with a Yokogawa CSU-X1 spinning disk; A 60x/ 1.42 NA oil Olympus objective was used together with 491 nm (100 mW; Cobolt), 561 nm (100 mW; Cobolt) and 645 nm (500 mW; Melles Griot) laser. A quad-edge dichroic beam splitter (446/ 523/ 600/ 677 nm; Semrock) was used to separate fluorescence emission from excitation light, and final images were taken with an sCMOS camera (Hamamatsu). Images were deconvolved using Huygens essential software (Scientific Volume Imaging).

STED imaging was performed using a custom designed Abberior775 3D-2 Color-STED system with 60x/1.2 NA Olympus water and 100x/1.4 NA oil Olympus objective. AlexFluor 594 was imaged with a pulsed laser at 560 nm, and excitation of Abberior Star Red was performed at 640 nm. The depletion laser for both colors was a Katana 775 nm pulsed laser. To reduce high frequency noise STED images were filtered with 2D or 3D Gaussian with a sigma of 0.8 pixels. The images were analyzed using Matlab (Mathworks) and Fiji^33^. Line profiles perpendicular to microtubules were Gaussian fitted to calculate the FWHM and average intensity. Distortion levels were estimated using the same scenario described before^1^. In brief, by applying a smoothing step based on anisotropic partial differential equation^34^, the pre-and post-expansion images were first smoothed and then fed into a non-rigid registration technique based on B-spline transformation model. Finally, the root mean square error between the pre-expansion image and the registered version of the post-expansion image were calculated using the technique suggested in and plotted as a curve.

## Acknowledgements

The authors thank J. Fuchs from B. Eickholt lab (Charité – Universitätsmedizin Berlin) for the cultured neurons, M. Rasband (Baylor College of Medicine) for the spectrin betaIV antibody, C. Martin-Lemaitre for preparing fixed samples, E. Boyden for help with data analysis, G. Pigino and G. Alvarez Viar for providing Chlamydomonas cells. This work was funded by DFG through SFB958 INST 130/827-2 to H.E., TRR186 INST 35/1409-1 to H.E and A.Z., TRR83 TP26 to A.H., SFB958 TP A16 and TRR186 TP A10 to B.E. and Freie Universität Berlin.

The authors acknowledge support from all Ewers lab members and Honigmann lab members for help and discussions.

## ORCID

HE: 0000-0003-3948-4332

## Author contributions

HE and AH designed research; MG, RM and OB performed research; BE contributed neuronal cultures; HE, AZ, MG analyzed data; MG, HE and AH wrote the paper with input from all authors

The authors declare no conflict of interest.

